# KDM6A facilitates Xist upregulation at the onset of X inactivation

**DOI:** 10.1101/2023.08.16.553617

**Authors:** Josephine Lin, Jinli Zhang, Li Ma, He Fang, Rui Ma, Camille Groneck, Galina N. Filippova, Xinxian Deng, Wenxiu Ma, Christine M. Disteche, Joel B. Berletch

## Abstract

X chromosome inactivation (XCI) is a female-specific process in which one X chromosome is silenced to balance X-linked gene expression between the sexes. XCI is initiated in early development by upregulation of the lncRNA *Xist* on the future inactive X (Xi). A subset of X-linked genes escape silencing and thus have higher expression in females, suggesting female-specific functions. One of these genes is the highly conserved gene *Kdm6a*, which encodes a histone demethylase that removes methyl groups at H3K27 to facilitate gene expression. Here, we investigate the role of KDM6A in the regulation of *Xist*. We observed impaired upregulation of *Xist* during early stages of differentiation in hybrid mouse ES cells following CRISPR/Cas9 knockout of *Kdm6a*. This is associated with reduced *Xist* RNA coating of the Xi, suggesting diminished XCI potency. Indeed, *Kdm6a* knockout results in aberrant overexpression of genes from the Xi after differentiation. KDM6A binds to the *Xist* promoter and knockout cells show an increase in H3K27me3 at *Xist*. These results indicate that KDM6A plays a role in the initiation of XCI through histone demethylase-dependent activation of *Xist* during early differentiation.

## Introduction

In mammalian females X chromosome inactivation (XCI) serves to balance X-linked gene expression between males (XY) and females (XX) [1, 2]. XCI is a robust silencing mechanism characterized by several chromosome-wide epigenetic changes on the Xi, including histone de-acetylation, ubiquitination of H2AK119, methylation of H3K27 and H3K9, DNA methylation at CpG islands, and chromatin condensation. XCI is initiated at early stages of development by upregulation of the long non-coding RNA (lncRNA) *Xist* from the future inactive X chromosome (Xi). *Xist* RNA coats the Xi and recruits proteins that implement epigenetic changes for silencing. This process can be modeled in mouse embryonic stem (ES) cells, which demonstrate extensive silencing by day 4-7 of differentiation [3, 4].

A subset of X-linked genes escape XCI and remain expressed from the Xi, albeit at a lower level than from the active X chromosome (Xa), resulting in higher expression in females compared to males [5, 6]. XCI and escape are processes unique to females, suggesting that factors involved in these processes such as *Xist* and other regulatory elements that influence the structure and epigenetic features of the Xi may be partly controlled by genes with female-biased expression [7]. One of these genes is *Kdm6a*, a highly conserved gene that escapes XCI in all mammalian species tested [8, 9]. The main function of KDM6A is to facilitate gene expression during differentiation and development by resolution of bivalent domains through demethylation of H3K27me3 [10-13]. Additionally, KDM6A plays a role in allelic gene regulation, where it preferentially targets maternal alleles of autosomal genes to increase expression [14]. Interestingly, KDM6A has a Y-encoded homolog (UTY) that has little or no demethylase activity, suggesting that KDM6A may have female-specific roles that depend on its demethylase function [15-17].

Here, we show that KDM6A plays a role in *Xist* upregulation during the earliest stages of XCI. We find that *Kdm6a* KO results decreased levels of *Xist* RNA and in reduced formation of an *Xist* cloud during ES cell differentiation. This causes impaired silencing of genes from the Xi. A direct association between H3K27me3 demethylation by KDM6A and *Xist* expression is supported by findings that KDM6A binds to the *Xist* promoter region during ES cell differentiation, and that *Kdm6a* KO results in an increase of H3K27me3 levels along the *Xist* gene. Thus, the female-biased expression of *Kdm6a* via escape from XCI may have evolved as a mechanism that contributes to XCI by regulating *Xist*.

## Results

### Generation of Kdm6a KO ES cells

To study the potential role of KDM6A at the onset of XCI, we used CRISPR/Cas9 to knockout (KO) *Kdm6a* in hybrid female mouse ES cells derived from a 129 x *Mus castaneus* (*cast*) cross in which alleles can be distinguished by SNPs (Figure S1A). In order to examine changes specifically on the Xi we used *Tsix-*stop cells (donated by J. Gribnau, Erasmus MC) in which a transcriptional stop signal is inserted onto the 129 allele of *Tsix*, resulting in completely skewed silencing of the 129 X chromosome upon differentiation [18]. Stable *Kdm6a* KO *Tsix-*stop ES cell clones with a homozygous deletion of part of exon 2 through part of exon 4 (ΔE/ΔE) were generated (Figure S1A) as previously derived [14]. PCR and Sanger sequencing verified homozygous editing of *Kdm6a* in two KO clones (*Tsix-Kdm6a*^*ΔE/ΔE17*^, *Tsix-Kdm6a*^*ΔE/ΔE21*^). Despite residual *Kdm6a* expression in the KO clones, there was no evidence of KDM6A protein [14, 15]. We also isolated control *Tsix-*stop ES cell clones (hereafter called CRISPR controls), which were subject to the CRISPR/Cas9 treatment but did not exhibit a deletion. In a separate experiment we derived clones E8-*Kdm6a*^*ΔP/ΔP26*^ and E8-*Kdm6a*^*ΔP/ΔP13*^, from another hybrid female mouse ES cell line called E8 (donated by J. Gribnau, Erasmus MC) (Figure S1A). In this cell line derived from a C57BL/6J (BL6) x *cast* cross, XCI is random, with partial skewing due to a strong Xce allele that results in a ∼3:1 inactivation of the BL6 X chromosome upon differentiation [19]. A summary of the clones used in this study can be found in Table S1.

### Kdm6a KO results in expression changes of genes involved in germ layer development

RNA-seq analyses were done to compare gene expression between wild-type (wt) and *Kdm6a* KO clones before (day 0, thereafter abbreviated D0) and after differentiation (D15) of ES cells into embryoid bodies (EB). For *Tsix-* stop cells, principal component analysis (PCA) based on either autosomal or X-linked gene expression shows little separation between wt and KO clones prior to differentiation, but a clear separation at D15, indicating that the effects of *Kdm6a* KO are most pronounced after differentiation (Figure 1A). Consistent with PCA clustering, we found only 703 differentially expressed genes (DEGs) between wt and KO cells at D0, while there were 1811 DEGs at D15 (≥2 fold change; FDR ≤0.1) (Figure S1B; Table S2). These findings are in agreement with KDM6A’s dispensable role in undifferentiated ES cells and its roles in regulating expression of genes involved in development and differentiation [11, 14, 20]. Indeed, among the DEGs found at D15 were a subset of genes involved in germ layer development. A subset of marker genes expressed in all three germ layers were downregulated in *Tsix-*stop *Kdm6a* KO cells at D15, but not at D0 prior to differentiation (Figure S1C; Table S3). Few changes were seen in E8 cells at D4 of differentiation (Figure S1C).

**Figure 1.**
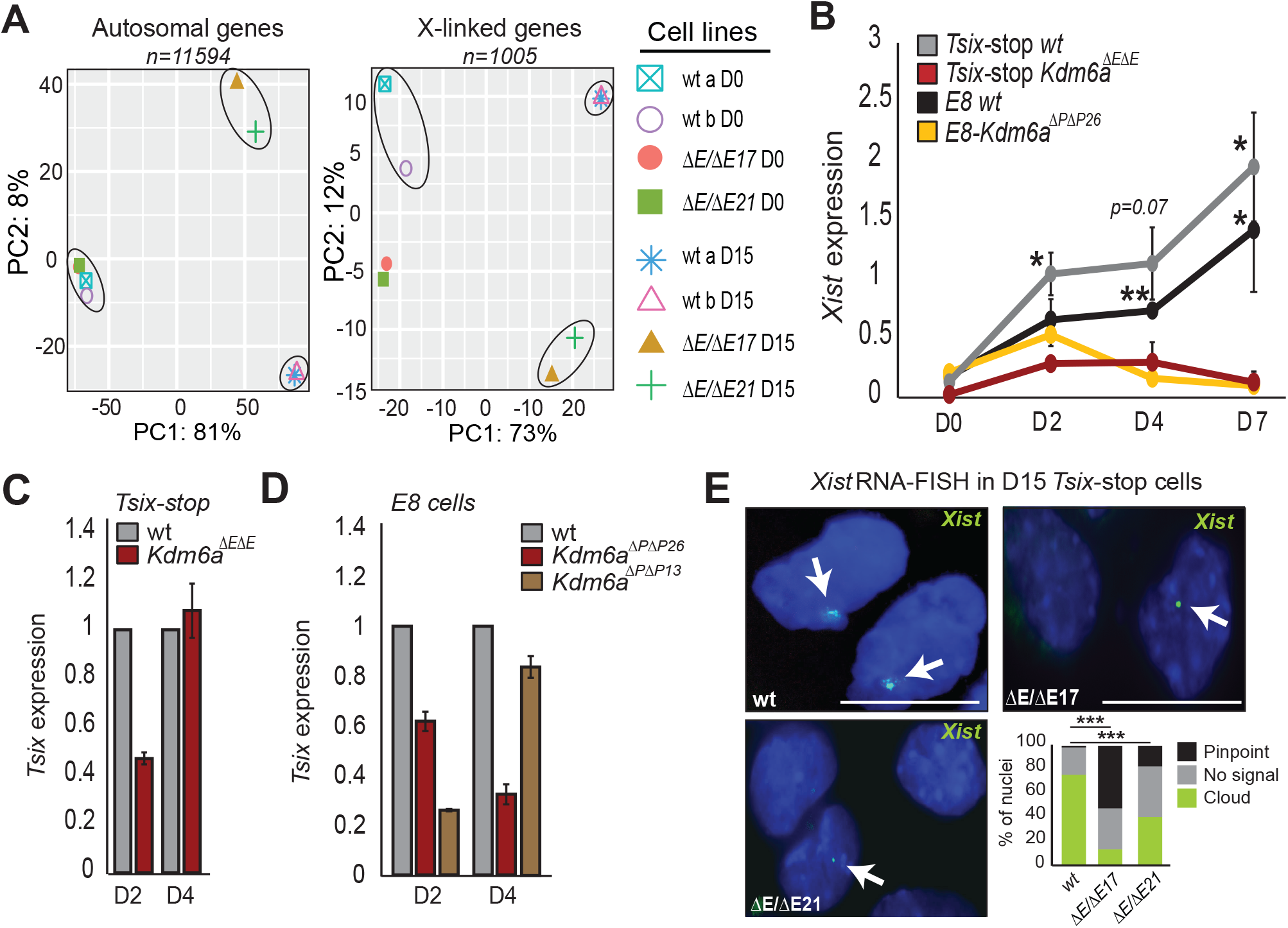
*Kdm6a* KO results in *Xist* downregulation. **(A)** Principal component analysis (PCA) based on expression of all autosomal or X-linked genes in two *Tsix-*stop wt clones and two *Kdm6a* KO clones (*Tsix-Kdm6a*^*ΔEΔE17*^, *Tsix-Kdm6a*^*ΔEΔE21*^) before (D0) and after differentiation (D15). Only genes with a minimum of 2.5 CPM in 4 libraries were considered. PCA shows separation of clones based on *Kdm6a* KO status mainly after differentiation. PCA figures were generated by iDEP.95. **(B)** qRT-PCR of *Xist* expression normalized to *Actinβ* during ES differentiation in *Tsix-*stop wt and *Kdm6a*^*ΔEΔE*^ KO clones (average values for *Tsix-Kdm6a*^*ΔEΔE17*^ and *Tsix-Kdm6a*^*ΔEΔE21*^), and in E8 wt and E8-*Kdm6a*^*ΔPΔP26*^ KO clone. p-values are derived from two biological replicates (**p<0.01; *p<0.05). Divergence of *Xist* expression between wt and KO cells begins at D2 and is largest at D7. **(C, D)** qRT-PCR of *Tsix* expression in **(C)** *Tsix-*stop wt and *Kdm6a*^*ΔEΔE*^ KO clones (see **B**), and (**D**) E8 wt and KO clones E8-*Kdm6a*^*ΔP/ΔP26*^ and E8-*Kdm6a*^*ΔP/ΔP13*^ at D2 and D4. Expression is normalized to *Actinβ* and relative to wt. **(E)** RNA-FISH with a probe specific for *Xist* RNA labeled in green in *Tsix-*stop wt and KO clones (*Tsix-Kdm6a*^*ΔEΔE17*^, *Tsix-Kdm6a*^*ΔEΔE21*^). Examples of nuclei are shown together with histograms for quantification of signal type. *Xist* clouds present in 74% of D15 wt cells are significantly decreased in KO clones (cloud compared to no signal; p<0.00001 for *Tsix-Kdm6a*^*ΔEΔE17*^ and <0.0001 for *Tsix-Kdm6a*^*ΔEΔE21*^ fisher’s exact test), while the frequency of pinpoint *Xist* signals increases in KO clones (pinpoint in wt compared to pinpoint in KO; p<0.00001 for *Tsix-Kdm6a*^*ΔEΔE17*^ and *Tsix-Kdm6a*^*ΔEΔE21*^ fisher’s exact test). Nuclei are counterstained with Hoechst 33342 Scale bar = 10μm.

### Xist expression and coating are impaired during early differentiation following Kdm6a KO

One of the earliest events at the onset of XCI is the upregulation of *Xist* [1, 2]. Therefore, we investigated changes in the dynamics of *Xist* expression during early mouse ES cell differentiation at D0, D2, D4, D7, and D15 following *Kdm6a* KO. In *Tsix-*stop cells we found that, in contrast to wt and CRISPR-control clones, *Kdm6a* KO clones (*Tsix-Kdm6a*^*ΔE/ΔE17*^, *Tsix-Kdm6a*^*ΔE/ΔE21*^) have significantly reduced levels of *Xist* expression starting at D2 of differentiation when *Xist* upregulation normally occurs (Figures 1B and S2A). This finding was confirmed in two *Kdm6a* KO clones (E8*Kdm6a*^*ΔP/ΔP13*^, E8-*Kdm6a*^*ΔE/ΔP26*^) obtained by editing ES line E8 (Figures 1B and S2B). Our observation that divergent expression patterns of *Xist* between wt and *Kdm6a* KO cells begin at D2 and become more obvious by D4 and D7 (Figure 1B) is consistent with published data from single-cell analysis [4]. Importantly, lower expression of *Xist* in *Kdm6a* KO cells does not seem to be due to changes of known *Xist* repressors or activators (*Tsix, Nanog, Pou5f1, Sox2, Rex1, Yy1, Rlim*), as these were not differentially expressed between wt and KO cells at D0 (Figure S2C) [21]. In particular, *Tsix* expression was consistently lower in KO cells at D2, reaching levels similar to wt only at D4, suggesting that the small increase in *Klf4* in KO cells did not affect *Tsix* expression [22] at this stage (Figures 1C, D and S2C; Table S2). Previous studies have shown aberrantly increased expression of *Xis*t with *Tsix* inhibition in undifferentiated ES cells grown in media with serum [23, 24]. However, the *Tsix-*stop ES cells used here maintain a low level of *Xist* expression at D0 similar to levels observed in previous studies despite of the presence of serum in the media (Figure S2D) [3, 14].

To determine whether *Kdm6a* KO affects the formation of an *Xist* cloud we performed RNA-FISH with a probe that targets *Xist* in differentiated *Tsix-*stop *Kdm6a* KO and wt cells at D15. The percentage of nuclei with *Xist* clouds sharply decreased after KO, with only 18% and 38% of nuclei with an *Xist* cloud in clones *Tsix-Kdm6a*^*ΔE/ΔE17*^ and *Tsix-Kdm6a*^*ΔE/ΔE21*^, respectively, versus 74% of wt nuclei (Figure 1E). Consistent with reduced *Xist* expression in *Kdm6a* KO clones, nuclei with a small pinpoint *Xist* signal rather than a cloud were prevalent in these clones (41% and 23% of nuclei in clones *Tsix*-*Kdm6a*^*ΔE/ΔE17*^ and *Tsix-Kdm6a*^*ΔE/ΔE21*^, respectively) compared to 4% in wt nuclei (Figure 1E). Loss of an X chromosome during cell culture is a common occurrence in female mouse ES cells and would confound our analysis of X-linked gene expression. To address this possibility DNA-FISH using probes targeting X chromosome-specific regions was done, which showed that little X chromosome loss occurred in any condition tested (Figure S3).

Together, these results indicate that KDM6A contributes to upregulation of *Xist* expression and to Xist coating of the Xi during early mouse ES cell differentiation.

### Aberrantly increased gene expression from the Xi in differentiated cells after Kdm6a KO

We postulated that reduced *Xist* upregulation and *Xist* coating in *Kdm6a* KO cells may lead to impaired XCI in these cells. Indeed, by RNA-seq the X:A expression ratio (ratio between the median expression of all X-linked genes versus all autosomal genes) was significantly increased in *Tsix-*stop *Kdm6a* KO (average of clones *Tsix-Kdm6a*^*ΔE/ΔE17*^ and *Tsix-Kdm6a*^*ΔE/ΔE21*^) compared to wt at D15 (Figure 2A). Allelic analyses of gene expression showed this increase in the X:A ratio to be due mainly to increased gene expression specifically from the Xi where 52% (225/431) of assayed genes show a ≥1.25 fold increase (p=0.002) (Figure 2B; Table S4). In contrast, there is a modest decrease of gene expression from the Xa (20% genes, p=0.28) and from the autosomes (28% genes, p=5.08^e-16^), consistent with the role of KDM6A in facilitating gene expression through removal of H3K27me3 at expressed genes (Figures 2B, C). This decrease in autosomal expression would contribute to a small extent to the increase in X:A ratio. An overall increase in X-linked gene expression was confirmed in an independent *Kdm6a* KO clone (E8-*Kdm6a*^*ΔP/ΔP13*^) where 43% (217/510) of X-linked genes show a ≥1.25 fold increased expression (p=0.07) by D4 of differentiation (Figure 2D; Table S4). Note that it was not possible to carry out allelic analysis in the E8 cell line due to random XCI albeit with skewing towards silencing of the B6 allele.

**Figure 2.**
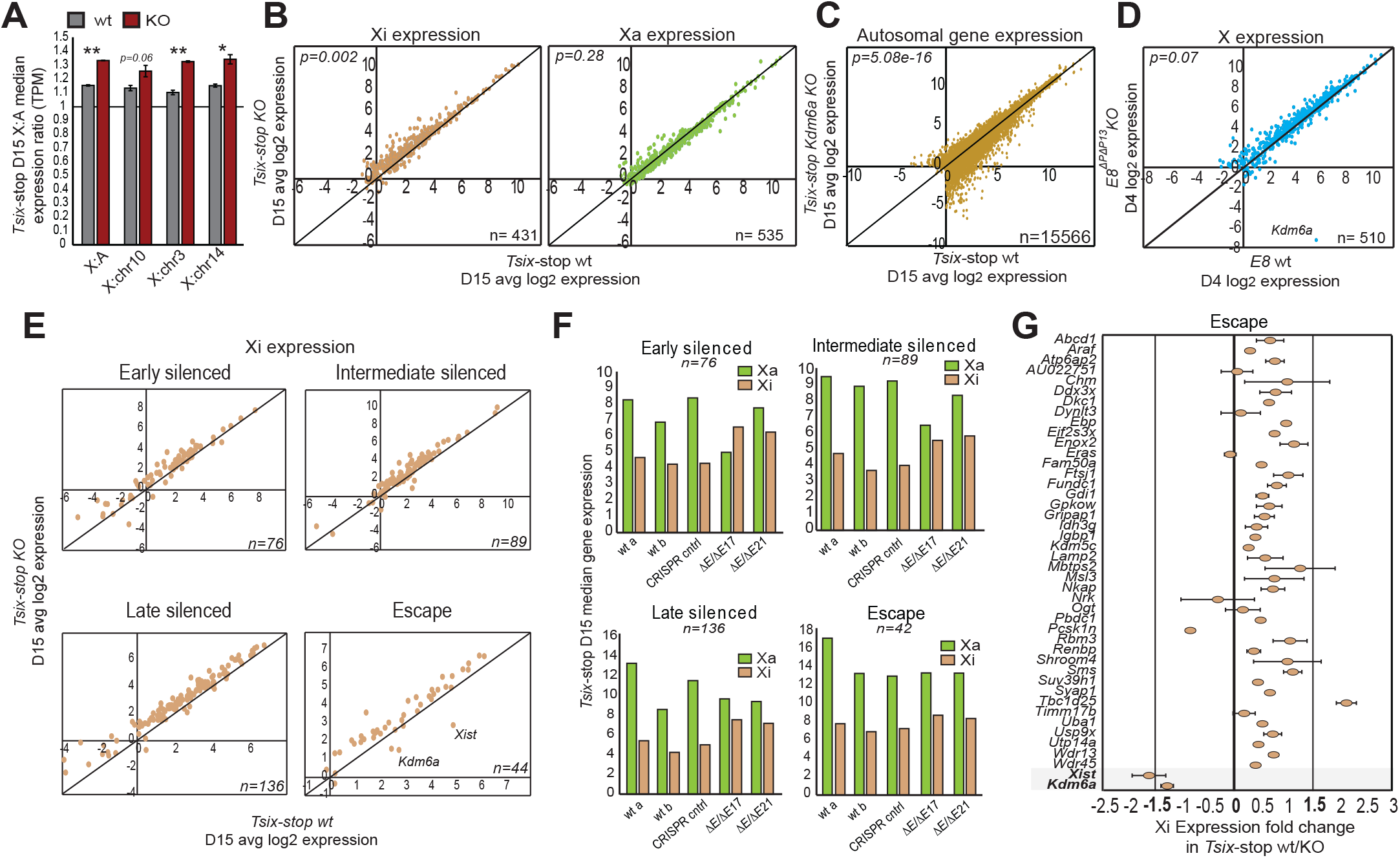
*Kdm6a* KO leads to increased gene expression from the Xi in differentiated cells. **(A)** Average X:autosome gene expression (X:A) ratio in *Tsix-*stop wt and KO clones (average for *Tsix-Kdm6a*^*ΔEΔE17*^ and *Tsix-Kdm6a*^*ΔEΔE21*^) at D15. Ratios between expressed X-linked and autosomal genes (>1TPM in a least two samples) are shown. Also shown are ratios between expressed X-linked genes and genes on three autosomes (chr10, chr3, and chr14) with a similar gene content as the X chromosome. There is increased X-linked gene expression following *Kdm6a* KO and differentiation. **(B)** Scatter plots of allelic expression from the Xi and Xa in *Tsix-*stop wt and KO clones (average for *Tsix-Kdm6a*^*ΔEΔE17*^ and *Tsix-Kdm6a*^*ΔEΔE21*^) at D15. Gene expression from the Xi but not the Xa is increased following KO. P-values are calculated using 1-way ANOVA test. n indicates the number of genes included. **(C)** Scatter plot of diploid autosomal gene expression in *Tsix*-stop wt and KO clones (average for *Tsix-Kdm6a*^*ΔEΔE17*^ and *Tsix-Kdm6a*^*ΔEΔE21*^) at D15 shows some decrease in expression in KO cells. Only genes with ≥1TPM in half the samples are shown. n indicates the number of genes included. **(D)** Scatter plot of total X-linked gene expression in E8 wt and KO clone E8-*Kdm6a*^*ΔP/ΔP13*^ at D4. Consistent with promoter KO, *Kdm6a* expression is abolished. There is a trend of higher expression of X-linked genes in KO cells. P-values calculated using 1-way ANOVA test. n indicates the number of genes included. **(E)** Scatter plots of expression between *Tsix-*stop wt and KO clones (average for *Tsix-Kdm6a*^*ΔEΔE17*^ and *Tsix-Kdm6a*^*ΔEΔE21*^) at D15 for X-linked genes categorized as those silenced early, silenced at an intermediate time, silenced late, and not silenced (escape) during XCI [3, 25]. KO clones show higher expression of genes in each category compared to wt. n indicates the number of genes in each category. **(F)** The median allelic expression of genes silenced at different times during XCI and of genes that escape XCI is increased in *Tsix-*stop KO clones compared to wt and CRISPR controls for all categories, while there is little change in expression from the Xa. n indicates the number of genes in each category. **(G)** Expression fold changes of individual escape genes from the Xi in *Tsix-*stop KO clones (average for *Tsix-Kdm6a*^*ΔEΔE17*^ and *Tsix-Kdm6a*^*ΔEΔE21*^) compared to wt at D15. Most genes show increased expression.

Next, we grouped X-linked genes in four defined temporal clusters, early, intermediate, late, and not silenced (or escape), based on their previously reported timing of silencing during differentiation [3, 25]. Genes classified as silenced early, intermediate, and late all show a trend of increased expression from the Xi in *Tsix-* stop *Kdm6a* KO cells compared to wt or CRISPR controls (Figures 2E, F; Table S5). Interestingly, the expression of most escape genes is also increased after *Kdm6a* KO (Figure 2E-G). The increase is usually less than 1.25 fold, with only three escape genes showing expression changes >1.25 fold in KO versus wt ES cells, including *Xist* and *Kdm6a*, both significantly decreased as expected, and *Tbc1d25*, which is increased (Figure 2G).

Taken together, these results indicate a reduced potency of XCI in differentiated *Kdm6a* KO cells, which implicates KDM6A, at least in part, as a regulator of X-linked gene expression dosage via upregulation of *Xist* during the onset of XCI.

### KDM6A binds to the Xist promoter

To explore the mechanisms of action of KDM6A in regulation of *Xist* we investigated whether KDM6A occupies the *Xist* locus during differentiation. Chromatin analysis using Cut&Run with an antibody for KDM6A in wt *Tsix-* stop cells at D2 and D7 shows that KDM6A binding to the *Xist* promoter region increases at D7, when *Xist* expression levels are most divergent between wt and KO cells (Figures 1B and 3A-C). Specifically, the fold change in *Xist* expression between D2 and D7 in wt *Tsix-*stop cells is consistent with the observed difference in KDM6A binding at the *Xist* promoter region at these time points (Figures 3B, C). Additionally, we investigated KDM6A binding at the *Dxz4* locus, which is known to play a role in maintenance of the condensed bipartite structure of the Xi [26-28]. Similar to our observations at the *Xist* locus, KDM6A binding shows a consistent increase across the *Dxz4* locus during differentiation (Figure S4).

**Figure 3.**
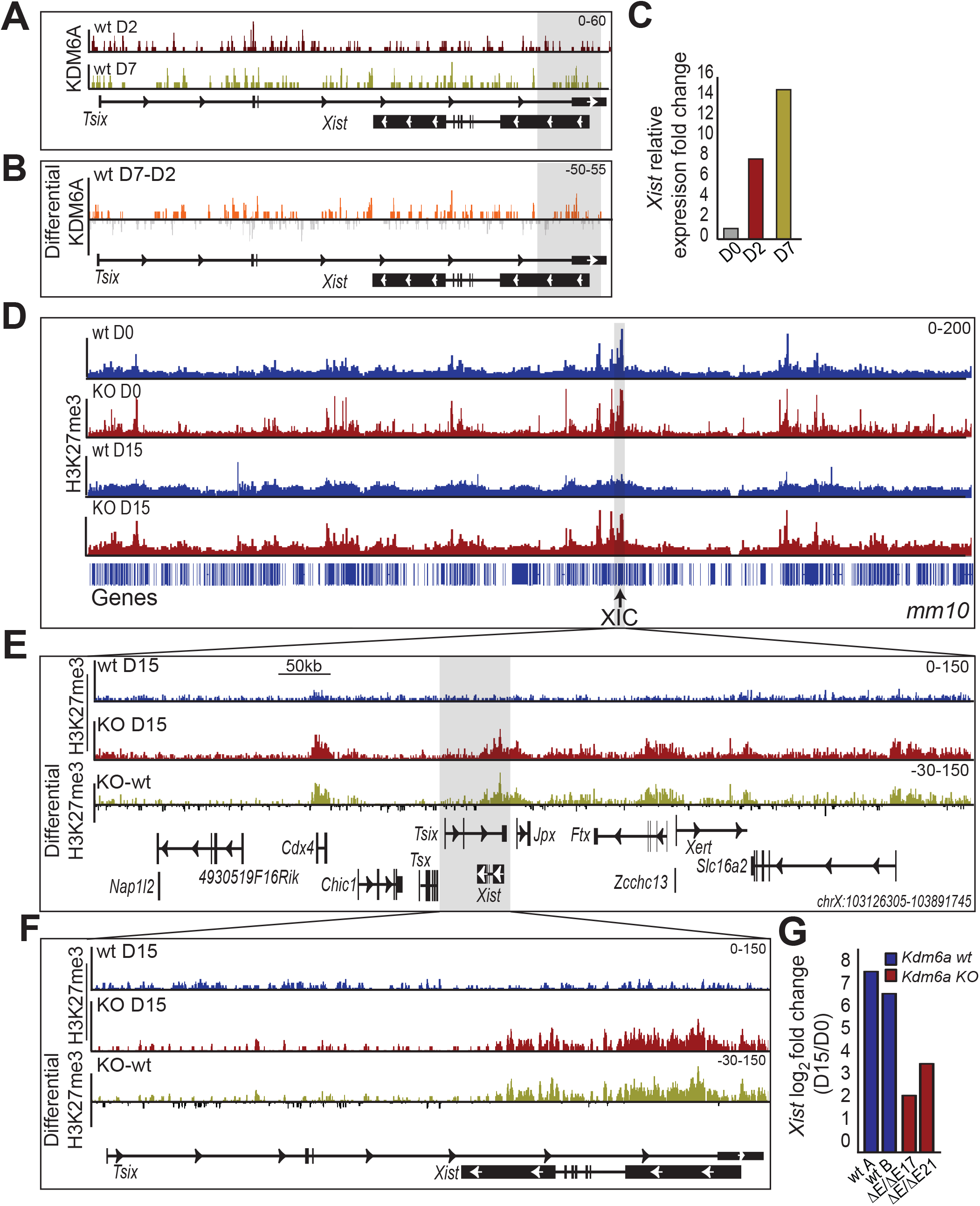
KDM6A binds to the promoter of *Xist* and *Kdm6a* KO results in gain of H3K27me3. **(A)** IGV browser view of KDM6A profile at D2 and D7 of differentiation at the *Tsix/Xist* locus in wt *Tsix*-stop cells. KDM6A occupancy at the *Xist* gene promoter region is modestly increased at D7. **(B)** Differential profile of KDM6A occupancy between D7 and D2 at the *Tsix/Xist* locus. Orange and gray signals indicate a gain and a loss in KDM6A occupancy, respectively. **(C)** *Xist* expression at D2 and D7 relative to D0 in *Tsix-*stop clones (average for *Tsix-Kdm6a*^*ΔEΔE17*^ and *Tsix-Kdm6a*^*ΔEΔE21*^) measured by qRT-PCR. Increased *Xis*t expression between D2 and D7 correlates with the increased KDM6A occupancy shown in **(A)** and **(B). (D)** IGV browser view of H3K27me3 profiles across the entire X chromosome in wt *Tsix*-stop cells (blue) and KO cells (*Tsix-Kdm6a*^*ΔEΔE17*^) (red) at D0 and D15. Blue bars underneath represent genes along the X chromosome. The XIC location is marked. **(E)** IGV browser view of H3K72me3 profiles at the XIC (chrX: 103126305-103891745) in wt *Tsix*-stop (blue) and KO *Tsix-Kdm6a*^*ΔEΔE17*^ cells (red). A differential profile between KO and wt is shown with yellow signals and black signals representing a gain and a loss in H3K27me3, respectively. **(F)** Same analysis as in **(E)** but zoomed in on the *Xist/Tsix* locus. **(G)** Log_2_ fold change of *Xist* expression in wt (2 isolates: wt A, wt B) and two KO clones (*Tsix-Kdm6a*^*ΔEΔE17*^ and *Tsix-Kdm6a*^*ΔEΔE21*^) at D15 relative to D0. There is a markedly higher level of *Xist* induction in wt compared to KO clones. The scales of the profiles shown in **(A), (B), (D), (E)**, and **(F)** are indicated in the upper right corners.

Our findings indicate that *Xist* and *Dxz4* bind KDM6A in wt ES cells. This may explain the increase in gene expression from the Xi we observed in *Kdm6a* KO cells, due to abnormal retention of H3K27me3 at loci critical for Xi silencing and maintenance of condensation, as shown below.

### Kdm6a KO induces an increase in H3K27me3 on the X chromosome

Next, we investigated changes in H3K27me3 profiles using Cut&Run to compare wt *Tsix-*stop cells to *Kdm6a* KO clone *Tsix-Kdm6a*^*ΔE/ΔE17*^. At D0, H3K27me3 levels along the whole X chromosome are similar between wt and KO cells, while at D15, most of the X chromosome demonstrates an increase in H3K27me3 levels in both wt and KO cells, as expected due to the onset of XCI (Figures 3D and S5A, B). However, this increase is more pronounced in *Kdm6a* KO compared to wt cells at D15 (Figure 3D). Note that wt cells at D15 show a decrease in H3K27me3 enrichment at the XIC, whereas high H3K27me3 levels persist at the XIC in KO cells. This is consistent with our findings of KDM6A binding to the *Xist* promoter region, which would remove methylation at H3K27 in wt cells to facilitate expression, but would fail to do so in KO cells (Figures 3E, F). The higher level of H3K27me3 at *Xist* in KO cells correlates with lower *Xist* expression, which culminates at only ∼2.5 fold upregulation between D0 and D15 in KO cells, compared to ∼6.5 fold in wt cells (Figure 3G). A closer inspection of the XIC reveals that H3K27me3 enrichment is markedly higher at the *Cdx4* gene, the *Tsix/Xist* locus, and the *Ftx* 5’ region in *Kdm6a* KO versus wt cells at D15 (Figures 3E and S5A). Both *Cdx4* and *Ftx* show little expression change following *Kdm6a* KO, however, these genes are expressed at very low levels following differentiation (<1TPM), precluding them from being classified as DEGs (Figures S5C-F). As a control, we analyzed a subset of known KDM6A target genes where increased H3K27me3 levels together with decreased expression were observed in KO versus wt cells (Figure S6A). Consistent with our previous study, the imprinted *Dlk1/Meg3* locus also showed increased H3K27me3 in *Kdm6a* KO cells. This increase, specifically located at the imprinting control regions (ICR), was concordant with decreased gene expression (Figure S6B; Table S2) [14].

Allelic effects of *Kdm6a* KO on the XIC were investigated in *Tsix-*stop cells in which there is maternally (129) skewed XCI [3]. At D0 few differences between alleles and between wt and *Kdm6a* KO are seen (Figures S7A, B). However, the future Xi (maternal X) shows enrichment in H3K27me3 along the *Xist* gene, which is not seen on the future Xa (paternal X) (Figure S7B; Table S6). This is evident both in wt and KO cells at D0, and is consistent with previous studies demonstrating maternal-specific H3K27me3 enrichment at *Xist* [29]. At D15, wt cells show high overall H3K27me3 levels on the 129 Xi compared to the *cast* Xa, as expected due to the onset of XCI (Figure 4A). *Kdm6a* KO results in even higher H3K27me3 levels on both the *cast* Xa and the 129 Xi (Figure 4A). Interestingly, only the maternal allele (129) of *Xist* shows persistence of higher levels of H3K27me3 following *Kdm6a* KO, which would explain decreased *Xist* expression from the 129 Xi and impaired XCI (Figures 4B, C; Table S6). Consistent with our diploid analysis above, allelic analyses show a marked increase in H3K27me3 on both alleles of *Cdx4* and *Ftx* at D15 (Figure 4B). A general increase in H3K27me3 may contribute to the large number of genes with decrease expression in *Kdm6a* KO cells versus wt (Figure S1B).

**Figure 4.**
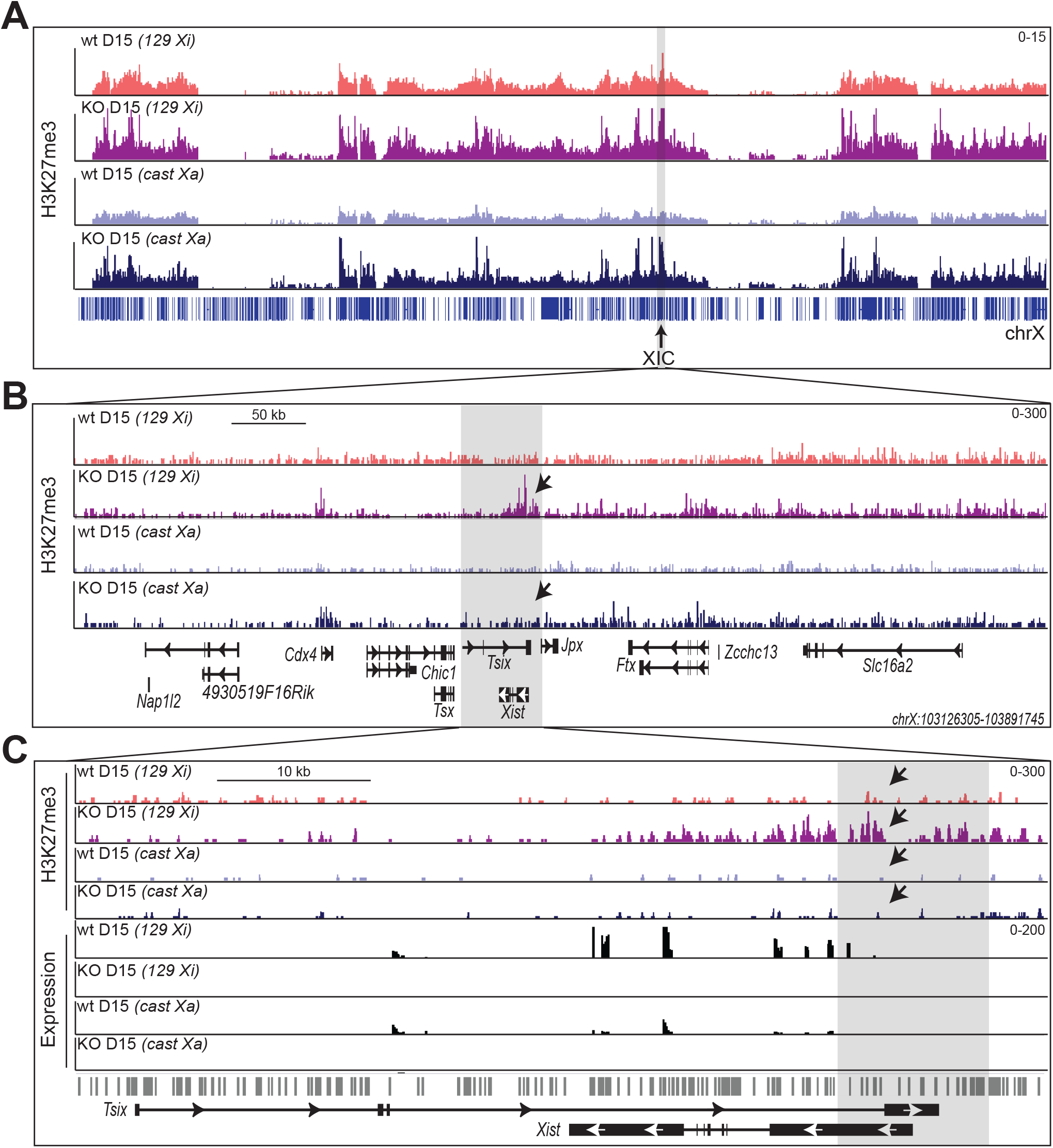
*Kdm6a* KO leads to increased H3K27me3 at *Xist* specifically on the Xi. **(A)** IGV browser views of H3K27me3 profiles across the entire Xi (129) and Xa (*cast*) obtained by allelic analysis in *Tsix-*stop wt and KO cells (*Tsix-Kdm6a*^*ΔEΔE17*^) at D0 and D15. The 129 Xi profile is in pink (wt) and purple (KO) and the *cast* Xa in light blue (wt) and dark blue (KO). Blue bars underneath represent genes along the X chromosome and the position of the XIC is marked. **(B)** Same analysis as in **(A)**, but zoomed in to show the XIC (chrX: 103126305-103891745). There is a marked increase in H3K27me3 at *Xist* on the Xi but not the Xa in KO cells (arrows). **(C)** Same analysis as in **(B)** but zoomed in on the *Xist/Tsix* locus. Also shown are allelic RNA-seq tracks of gene expression. The Xi-specific increase in H3K27me3 enrichment at the *Xist* promoter (arrow) is accompanied by lower *Xist* expression from the 129 Xi allele. No expression of *Xist* was observed from the cast allele, which is the Xa. SNPs are shown by gray lines below IGV tracks. The scales of the profiles are indicated in the upper right corners.

Our findings show that prior to the onset of XCI, *Kdm6a* KO has little effect on H3K27me3 levels at the XIC. Indeed, H3K27me3 levels remain high at *Xist* on the future Xi after KO in undifferentiated cells. Following differentiation persistence of high H3K27me3 levels is apparent in *Kdm6a* KO cells, concurrent to low levels of *Xist* expression. Although we cannot exclude indirect effects of *Kdm6a* KO on *Xis*t regulation, our data showing association of KDM6A with the 5’ end of *Xist* during differentiation suggests that KDM6A may directly facilitate *Xist* expression during XCI.

## Discussion

Our study implicates KDM6A in the upregulation of *Xist* at the onset of XCI. We find that the female-biased histone demethylase helps *Xist* upregulation by binding to its promoter region and demethylating H3K27me3 during the early days of mouse embryonic stem cell differentiation. In differentiated cells and adult tissues *Kdm6a* has higher expression in females compared to males due to escape from XCI, which may have evolved as a mechanism to regulate female-specific processes such as XCI [30]. Previous studies have shown that maternal imprinting of *Xist* is maintained by maternal-specific H3K27me3 at *Xist* in blastocysts and that demethylation is critical for initiation of *Xist* expression and XCI. Indeed, injection of *Kdm6b*, a gene that is located on chromosome 11 and encodes another H3K27me3 demethylase, in preimplantation embryos results in maternal allele-specific loss of H3K27me3 and induction of *Xist* expression [29]. Therefore, removal of the repressive chromatin mark appears necessary to initiate random XCI. Here, we show that KDM6A facilitates upregulation of *Xist* during early stages of XCI in hybrid mouse ES cell models with skewed or near-random XCI. *Kdm6a* KO leads to increased H3K27me3 levels on the maternal allele of *Xist* whose expression and ability to coat the inactive X decrease following differentiation. In turn, this failure of XCI results in aberrant expression of X-linked genes from the Xi. The ES cell line in which we followed changes in *Xist* expression and in silencing of X-linked genes has skewed XCI of the maternal 129 allele due to a stop mutation at *Tsix*. It could be argued that this mutation may affect our observations. However, we found that an independent *Kdm6a* KO line without a *Tsix* mutation also shows impaired *Xist* expression and X silencing.

We have previously demonstrated that KDM6A regulates expression of the X-linked *Rhox* genes in a female-specific manner via removal of H3K27me3 [31]. In addition, we showed that KDM6A preferentially regulates maternal alleles of autosomal genes [14, 31]. Transcriptional regulation by KDM6A can occur through histone demethylation-dependent and independent mechanisms [20]. In contrast, the protein encoded by the Y-linked homolog of *Kdm6a*, UTY, lacks detectable demethylase activity, suggesting that female-biased expression of *Kdm6a* may have evolved female-specific functions requiring H3K27me3 demethylation [15, 20]. Consistent with this, homozygous knockout (KO) of *Kdm6a* is lethal in female mice, while a small number of runted males with hemizygous deletion survive, suggesting that UTY partially compensates for loss of KDM6A through demethylation-independent mechanisms [32].

XCI is necessary for loss of pluripotency and normal differentiation, a mechanism that depends on expression of specific genes [4, 33, 34]. We find that germ layer marker genes have lower expression in *Kdm6a* KO cells than in wt, which is consistent with previous analyses demonstrating that KDM6A impairs differentiation. Indeed, *KDM6A* mutations are associated with increased “stemness” in cancer and *Kdm6a* KO impairs mesoderm development *in vivo* largely through histone demethylase-independent mechanisms [20]. However, recent studies of *Kdm6a* KO models indicate that depletion of the enzyme does not block differentiation, and may even facilitate differentiation into neuroectoderm lineages [35, 36]. Indeed, we only observed effects on germ layer marker genes after XCI has occurred and our KDM6A chromatin binding analyses suggest a direct effect of KDM6A on *Xist* expression during early XCI rather than an effect on differentiation itself. Allelic single cell RNA-seq analyses in differentiating hybrid ES cells show that XCI is complete prior to lineage specification, suggesting that the effects of *Kdm6a* KO on *Xist* we observed occur before lineage determination [4]. Thus, increased X-linked gene expression due to impaired XCI because of KDM6A deficiency may contribute to phenotypes in a variety of tissues such as those observed in Kabuki and Turner syndromes [37, 38]. However, our observation of increased expression of *Klf4* and other lineage-specific genes following *Kdm6a* KO suggests delayed differentiation, therefore, we cannot fully decouple the potential effects of *Kdm6a* KO on differentiation from the effects on regulation of *Xist* expression.

The *Xist* promoter is maintained in a bivalent state in undifferentiated ES cells and therefore is probably targeted for resolution by KDM6A via removal of H3K27me3 to facilitate expression during initiation of XCI [13, 39-41]. Consistent with this hypothesis, KDM6A occupancy appears to increase, albeit slightly, at the *Xist* promoter during early XCI in wt cells, coincident with increased *Xist* expression. KDM5C, which is also encoded by an escape gene, has recently been implicated in facilitating *Xist* expression by converting H3K4me3 to H3K4me1 at the *Xist* 5’ end [7]. Interestingly, KDM5C appears to be enriched at a similar location at the *Xist* gene as KDM6A, suggesting potential cooperation between escapees that may help form a permissive chromatin environment for expression.

Our results suggest a role for KDM6A in regulating the expression of regulatory elements located in the XIC, such as *Ftx* and *Jpx*, which have known roles in *Xist* regulation [42]. Thus, KDM6A may regulate *Xist* via direct or indirect mechanisms. *Ftx* transcription promotes *Xist* activation during the initiation of XCI [43], and it is possible that, similar to *Xist, Ftx* transcription is impaired during differentiation of *Kdm6a* KO cells. While *Ftx* has very low expression in our *Tsix-*stop ES cells (<1TPM), we did observe an increase of H3K27me3 levels at the *Ftx* promoter and a slight decrease in expression following *Kdm6a* KO. Notably, lower expression of *Tsix* was observed during early differentiation in *Kdm6a* KO cells compared to wt, indicating that the decrease in *Xist* expression we observe following KO is independent of *Tsix-*induced repression [44].

The polycomb complexes PRC1/2 that mediate methylation of H3K27 have been implicated in the dynamics of XCI and maternally imprinted genes [29, 45, 46]. Indeed, the absence of polycomb group proteins due to lack of *Xist*-mediated recruitment leads to a relaxation of transcriptional silencing [47]. Consistent with these results, we observe a significant upregulation of gene expression from the Xi following differentiation in *Kdm6a* KO cells. A trend of increased expression from the Xi was seen for all temporally defined clusters of silenced genes, as well as genes in the escape category. Genes that escape XCI rarely achieve a level of Xi expression equal to that from the Xa due the repressive chromatin environment of the Xi [37]. This suggests that effects of *Kdm6a* KO on genes that escape may be due to an overall relaxation of the Xi heterochromatin. Interestingly, we observed KDM6A occupancy at *Dxz4*, a locus known to play a role in the Xi 3D structure by maintaining the two condensed superdomains of the Xi [26-28]. KDM6A occupancy at *Dxz4* during differentiation suggests that it may play a regulatory role in the overall architecture of the Xi. However, further investigation is needed to determine what that role may be.

Here, we illustrate a novel female-specific role for the X-linked histone demethylase KDM6A in the regulation of X-linked gene expression dosage via its involvement in the initiation of XCI through its histone demethylation-dependent function. Further studies are needed to determine if the effects of KDM6A depletion on *Xist* are direct or indirect. A recent study points to a role for KDM6A in regulating *XIST* expression in human disease [48]. Therefore, extending our studies to human models will help determine the role of KDM6A in regulating X-linked gene expression in cases of sex chromosome aneuploidy as well as Kabuki syndrome and in cancer, where it is frequently mutated.

## Authors’ contributions

Conceptualization, J.B. and C.M.D.; Investigation, J.B., J.L., H.F., C.G.; Formal analysis, H.F., J.Z., L.M., R.M., W.M.; Data Curation, W.M.; Writing – Original Draft, J.B. and C.M.D.; Writing – Review and Editing, J.B., J.L., W.M., G.N.F., and C.M.D.; Visualization, J.B., W.M., X.D., and C.M.D.; Supervision, J.B. and C.M.D.; Funding Acquisition, J.B. and C.M.D.

## Acknowledgements

We thank Di Kim Nguyen (University of Washington) for consultation on this study and for critical reading of the manuscript and Charlie Lee (University of Washington) for his help with next-generation sequencing. Hybrid mouse ES cells used in this study were graciously donated by Joost Gribnau (Erasmus MC, University Medical Center).

## Experimental Methods

### Cell culture and differentiation

The F1 female mouse hybrid ES line *Tsix-*stop was derived from a cross between 129 and *cast* followed by the insertion of a transcriptional stop in the *Tsix* gene on the 129 X chromosome [18]. Differentiation of *Tsix-*stop ES cells results in skewed XCI where the Xi is always the 129 (maternal) X chromosome. In contrast, differentiation of the F1 female mouse hybrid ES line, E8, derived from a cross between C57BL/6J (B6) and *cast* results in random XCI. ES cells were maintained in the presence of 1000 U/ml leukemia inhibitory factor (LIF) (Millipore) on a monolayer of chemically inactivated mouse embryonic fibroblasts (MEFs), in high glucose DMEM media supplemented with 15% fetal bovine serum (FBS), 1% non-essential amino acids, 10 mg/ml APS, 0.1mM 2mercaptoethanol and 25mM L-glutamine, in a humidified incubator at 37ºC and 5% CO_2_. Just prior to use, ES cells were enriched by incubation on 0.1% gelatin-coated dishes for 1h to allow MEFs to attach, followed by transfer to fresh gelatin-coated plates for overnight culture. Following expansion, ES cells were split once (1:10) to further reduce potential MEF contamination. For embryoid body (EB) formation, 4×10^6^ wt control and *Kdm6a* KO cells were cultured on non-adherent bacterial culture dishes without LIF for 7 days. Cells were then plated onto gelatin-coated cell culture plates and maintained for another 8 days prior to collection at D15.

### Kdm6a editing using CRISPR/Cas9

We chose a dual sgRNA approach because of the capacity to create large deletions and reduced chance of off-target effects [49]. Editing of *Kdm6a* in *Tsix-*stop ES cells was done by constructing CRISPR plasmids containing sgRNAs that target exons 2 and 4 to delete ∼45kb, as done previously [14, 50]. ES cells were transfected with the CRISPR/Cas9 constructs and a plasmid carrying puromycin resistance using UltraCruz® transfection reagent at a 3:1 CRISPR to pPGKpuro ratio in media with no antibiotics. Two days later, cells were selected in ES media containing 1μg/ml puromycin for 48-72h, followed by recovery in media with antibiotic. Cells were then cloned into 96-well plates using serial dilutions. Clones were expanded and screened for deletions using PCR and Sanger sequencing to confirm the predicted homozygous ∼45 kb deletion, which was found in two *Tsix-*stop ES cell clones (*Tsix-Kdm6a*^*ΔE/ΔE17*^, *Tsix-Kdm6a*^*ΔE/ΔE21*^) (Figure S1A; Table S1). Additionally, we confirmed that neither *Tsix-Kdm6a*^*ΔE/ΔE1*^ nor *Tsix-Kdm6a*^*ΔE/ΔE2*^ CRISPR controls were edited (Figure S1A). Editing of *Kdm6a* in E8 ES cells was done by targeting the *Kdm6a* promoter region using sgRNAs designed using the CHOPCHOP v2 online tool [51]. PCR amplification and Sanger sequencing confirmed the predicted ∼5 kb homozygous promoter deletion in two E8 derived clones (E8-*Kdm6a*^*ΔE/ΔP26*^, E8-*Kdm6a*^*ΔP/ΔP13*^) (Figure S1A; Table S1). Plasmid preparations were made using either the Qiagen Mini-prep or Maxi-prep kits according to the manufacture protocol. Plasmid DNA and RNA lysates used for Sanger sequencing were prepared and submitted to Eurofin Genomics according to the protocols provided (http://www.operon.com)

### DNA and RNA isolation and qRT-PCR

RNA and DNA lysates were prepared using Qiagen RNeasy mini kit and the Qiagen DNeasy blood and tissue kit, respectively. cDNA synthesis was done with the GoScript reverse transcription kit (Promega). Relative transcript levels were determined using SYBR Green PCR master mix on an ABI 7900HT machine. qRT-PCR assays were conducted in triplicates and normalized to *Actinβ* prior to analysis using the comparative CT method.

### DNA-FISH and RNA-FISH

To verify the presence of two X chromosomes in differentiated wt and CRISPR treated cells, DNA-FISH was done on metaphase cells fixed in methanol:acetic acid (3:1 volume) using BAC probes (RP23-299L1 for *Dxz4* or RP24-322N20 for *Firre*) [52]. Probes were labeled with SpectrumGreen (Vysis #02N32-050) using a nicktranslation reagent kit according to the manufacturer’s protocol (Abbott Molecular Inc.). Probes in hybridization buffer were denatured at 70 °C for 5 min and hybridized overnight at 37 °C to slides that were also denatured at 70 °C for 5 min prior to hybridization. On the next day, slides were washed in SSC containing NP40 at 73 °C, followed by washes at room temperature. Counterstaining was done with Hoechst 33342 (2 μg/ml) in phosphate buffered saline (PBS) and slides were mounted in anti-fade solution.

For *Xist* RNA-FISH cells were plated on chamber slides at D7 of differentiation and allowed to differentiate until D15 when cells were processed directly on the chamber slides as described [53]. Briefly, cells were washed in PBS and permeabilized in CSK buffer with 0.05% Triton. Following fixation in 4% paraformaldehyde, cells were washed in 70% ethanol and dehydrated for 3 min each in 70%, 85%, and 100% ethanol. Dehydrated slides were incubated with a denatured probe for *Xist* RNA overnight in 50% formamide/50% 2xSSC at 42 °C in a humidified chamber [53]. On the following day, slides were washed three times in 50% formamide/50% 2xSSC at 42 °C for 10 min followed by one wash in 2xSSC at 42 °C for 5 min. Counterstaining was done with 33342 Hoechst in 2xSSC. A minimum of 130 nuclei were scored for each condition to quantify the number of nuclei with either an *Xist* cloud, or a pinpoint *Xist* signal, or no *Xist* signal.

### RNA-seq and allelic gene expression analysis

RNA-seq indexed libraries were prepared using Illumina TruSeq RNA sample preparation kit V2. For *Tsix-*stop cells, libraries were prepared from two biological replicates from wt controls, one CRISPR control (cln1) and *Tsix*-*Kdm6a*^*EΔ/EΔ17*^ and *Tsix*-*Kdm6a*^*EΔ/EΔ21*^ KO clones at D0 and D15. For E8 cells, libraries were prepared from one wt control and one E8-*Kdm6a*^*ΔP/ΔP13*^ clone at D4. Sequencing on a NextSeq sequencer yielded 75bp single-end reads. Diploid gene expression was estimated using Tophat/v2.0.14 [54] with default parameters and gene-level expression was normalized using TPM (transcripts per kilobase of exon length per million mapped reads). Differentially expressed genes were determined using DESeq2 analysis [55] with a false discovery rate (FDR) threshold of 0.1 and a fold-change cutoff of 2. PCA plots and heatmaps were generated with iDEP.95 using raw read counts as the input. Only genes with a minimum expression value of 2.5 CPM in at least two libraries were included (http://bioinformatics.sdstate.edu/idep95/).

To estimate allele-specific gene expression in mouse hybrid cells, the pseudo-diploid genome of 129x*cast* was generated using GATK-fastagenerator subcommand [56], taking the mm10 reference genome fasta file and the corresponding VCF file as the input. The VCF file was downloaded from the Sanger mouse genomes project [57], with indels and low-quality SNPs filtered out. RNA-seq reads were mapped using bowtie2 [58] to the diploid pseudo-129 and pseudo-*cast* genomes and transcriptomes. Only those reads that mapped uniquely and with a high-quality mapping score (MAPQ ≥ 30) were kept for further analyses. After filtering, reads were segregated into three categories: (1) 129-SNP reads containing only 129-specific SNPs; (2) *cast*-SNP reads containing only *cast*-specific SNPs; (3) allele-uncertain reads, that is, reads that do not contain valid SNPs. Allelic mapping metrics are shown in Table S7.

### Epigenetic analyses by CUT&RUN

Cut&Run for KDM6A was done in wt cells using an antibody against KDM6A (Cell signaling) according to the manufacturer’s protocol (EpiCypher). Cut&Run for H3K27me3 was performed on clone *Tsix-Kdm6a*^*ΔE/ΔE17*^ and wt cells using an antibody against H3K27me3 (Millipore) following a published protocol [59]. Libraries were constructed using TruSeq DNA sample preparation kit (Illumina) following the manufacturer protocol with minor changes, including two 1.1x bead purifications during the final selection step. Libraries were sequenced on a NextSeq sequencer to generate 75bp paired-end reads. In the following text, we refer to paired-end reads simply as “reads”. Demultiplexed fastq files were mapped to NCBi v38/mm10 mouse genome using bowtie2 [58] with the following parameters: `--end-to-end --very-sensitive --no-mixed --no-discordant --phred33 -I 10 -X 700`. Only paired reads that were mapped to the same chromosome with a fragment length of less than 1000bp were kept for further analyses. Read alignment BAM files were converted to bigWig format for IGV visualization using Bedtools [60]. Read counts were normalized using RPKM; and IgG control was subtracted from the knockout sample using the bamcompare subcommand in deeptools [61].

For allelic-analysis, the pseudo-diploid genome of 129 x *cast* was generated using GATK-fastagenerator subcommand [56], taking the mm10 reference genome fasta file and the corresponding VCF file as the input. The VCF file was downloaded from the Sanger mouse genomes project [57], with indels and low-quality SNPs filtered out. Reads were mapped to the diploid pseudo-129 and pseudo-*cast* genomes using bowtie2 [58] with the following parameters: `--end-to-end --very-sensitive --no-mixed --no-discordant -phred33 -I 10 -X 700`. Reads were further processed through a read tagging step, where read alignments of both pseudo-129 and pseudo-CAST genomes were compared to identify the allele origins of the reads. Reads overlapping SNPs were assigned to G1 (129 genome), G2 (*cast*), or CF (conflicting) based on the sequenced base information at SNP sites, while reads do not overlap with any SNPs were assigned to UA (Unassigned). The G1 and G2 tagged reads were separated into two BAM files, which were later used to generate allelespecific bigWig files for IGV visualization [62]. An allelic mapping summary can be found in Table S6.

## Supplemental Figure Legends

**Supplemental figure S1. CRISPR-Cas9 strategy and characteristics of *Kdm6a* KO cells. (A)** Top: Schematic shows the location of the exonic deletion (*Kdm6a*^*ΔE*^) that removed exons 2-4 of *Kdm6a* in female *Tsix-*stop ES cells and the promoter targeted deletion (*Kdm6a*^*ΔP*^) made in female E8 ES cells. Exons are shown as vertical bars. The location of the PCR and RT-PCR primers (color-coded arrows) used to confirm the deletion is indicated. Below left: Images of gels after electrophoresis of PCR products to confirm *Kdm6a* homozygous KO (-no deletion; + deletion positive). Below right: Partial *Kdm6a* sequence obtained by Sanger sequencing as verification of deletions in all four *Kdm6a* KO clones. Arrows point to the location of non-homologous end-joining and colored arrows correspond to those on the schematic. **(B)** Heat maps of gene expression differences between *Tsix-*stop wt (2 replicates, wt A, wt B) and KO clones (*Tsix-Kdm6a*^*ΔEΔE17*^ and *Tsix-Kdm6a*^*ΔEΔE21*^) at D0 and D15. Consistent with PCA clustering, more DEGs were identified at D15 than D0. Heat maps were generated using iDEP.95. **(C)** Scatter plots of expression of genes involved in germ layer differentiation obtained by RNA-seq in *Tsix-*stop and E8 wt and KO cells (*Tsix-Kdm6a*^*ΔEΔE17*^, *Tsix-Kdm6a*^*ΔEΔE21*^; E8-*Kdm6a*^*ΔPΔP13*^) during differentiation show higher expression of germ layer-associated genes in *Tsix*-stop wt cells at D15 and a trend of higher expression at D4 in E8 wt cells. For *Tsix-*stop, log_2_ average TPM values are from two wt replicates and two KO clones. For E8, log_2_ TPM values are from one wt and one KO clone. P-values are calculated using 1way ANOVA test.

**Supplemental figure S2. Confirmation of *Xist* expression changes in *Kdm6a* KO cells. (A)** qRT-PCR for *Xist* expression during differentiation in *Tsix-*stop wt and CRISPR controls (Table S1). Expression is normalized to *Actinβ*. Expression at D4 is relative to D0 for each cell line. As expected, *Xist* is upregulated upon differentiation in wt (wt A and wt B replicates) and CRISPR-control clones (cln1 and cln2). **(B)** qRT-PCR of *Xist* expression in E8 wt and an additional KO clone (E8-*Kdm6a*^*ΔPΔP13*^). Expression is normalized to *Actinβ. Xist* is significantly lower at D2 and D4 in KO cells. P values derived from two technical replicates of wt and KO cells (**p<0.01;*p<0.05). **(C)** Log_2_ of average TPM values for pluripotency genes and genes known to play a role in *Xist* repression. Average TPM values are from two *Tsix-*stop wt and two *Kdm6a* KO clones (*Tsix-Kdm6a*^*ΔEΔE17*^ and *Tsix-Kdm6a*^*ΔEΔE21*^). Only *Klf4* is called as differentially expressed between wt and KO by DESeq2 (**p<0.01). **(D)** Average TPM values for *Xist* in *Tsix-*stop wt, a CRISPR-control clone (cln1), and two KO clones (*Tsix-Kdm6a*^*ΔEΔE17*^, *Tsix-Kdm6a*^*ΔEΔE21*^) at D0, confirming that *Xist* expression is very low in ES cells cultured with serum. **Supplemental figure S3. Two X chromosomes are present in wt and KO *Tsix*-stop and E8 cells. (A, B)** DNA-FISH using probe*s* specific for the X-linked gene *Firre* or an X-linked mini-satellite repeat region (*Dxz4*) labelled in green in *Tsix*-stop wt **(A)** and CRISPR-control (cln1) **(B)**. Examples of nuclei with two green signals representing the two X chromosomes are shown along with histograms of the number of signals in nuclei scored. n indicates the number of nuclei scored. Nuclei are counterstained with Hoechst 33342. **(C, D)** Same analysis in as in **(A, B)**, but in the *Tsix*-stop *Kdm6a* KO cell clones (*Tsix-Kdm6a*^*ΔEΔE17*^, *Tsix-Kdm6a*^*ΔEΔE21*^). **(E, F)** Same analysis as in **(A, B)**, but in E8 wt and KO E8-*Kdm6a*^*ΔPΔP26*^ clone. All wt and CRISPR treated clones show two signals in 77***-***90% of cells, confirming retention of both X chromosomes.

**Supplemental figure S4. KDM6A occupies the *Dxz4* region in wt *Tsix-*stop cells**. IGV browser view of profiles of KDM6A binding at D2 (red), D4 (blue), and D7 (yellow) of differentiation at the *Dxz4* locus in wt *Tsix-* stop cells. KDM6A occupancy at the *Dxz4* locus modestly but consistently increases during differentiation. The scale of the profiles is indicated in the upper right corner.

**Supplemental figure S5. H3K27me3 profiles in wt and *Kdm6a KO* ES cells at D0. (A)** IGV browser view of profiles of H3K27me3 enrichment at the XIC (chrX: 103126305-103891745) in undifferentiated (D0) *Tsix-*stop wt (blue) and KO clone *Tsix-Kdm6a*^*ΔEΔE17*^ (red). **(B)** Same analysis as **(A)**, but zoomed in on the *Tsix/Xist* locus. Shown underneath in black is the RNA-seq track, which shows little to no *Xist* expression at D0. **(C, D)** Expression (TPM) of *Cdx4* **(C)** and *Ftx* **(D)** in undifferentiated *Tsix-*stop wt (blue) and *Kdm6a* KO clones (red). Both wt and KO ES cells show low expression of *Cdx4* and *Ftx* (∼1.5TPM or less). **(E, F)** Expression (TPM) of *Cdx4* **(E)** and *Ftx* **(F)** in differentiated *Tsix-*stop wt (blue) and *Kdm6a* KO clones (red). Both wt and KO ES cells show low expression of *Cdx4* and *Ftx* (∼1TPM or less). The scale of the profiles shown in **(A**) and **(B)** is indicated in the upper right corner.

**Supplemental figure S6. Epigenetic and expression changes at known KDM6A target genes. (A)** IGV browser views of profiles of H3K27me3 enrichment at known KDM6A target genes in differentiated *Tsix-*stop wt cells (blue) and KO clones (*Tsix-Kdm6a*^*ΔEΔE17*^) (red). Below are RNA-seq expression profiles (black) in wt and KO cells. Histograms show TPM expression values for *T* and *Pitx2* in two differentiated wt clones and KO clones. For *Wnt3* and *Gata4*, qRT-PCR values during differentiation and TPM values following differentiation are also shown (*p<0.05). KO represents an average for the *Tsix-Kdm6a*^*ΔEΔE17*^ and *Tsix-Kdm6a*^*ΔEΔE21*^ clones. Values are normalized to *Actinβ*. **(B)** Same analysis as in **(A)**, but at the *Dlk1*/*Meg3* imprinted locus at D15. H3K27me3 markedly increases at the promoter of *Meg3*, which is the imprinting control region ICR (2) as well as the intergenic ICR (1) in KO cells. The scales of the profiles shown in **(A)** and **(B)** are indicated in the upper right corners.

**Supplemental figure S7. H3K27me3 allelic profiles in wt and *Kdm6a KO* ES cells at D0. (A)** IGV browser view of H3K27me3 enrichment mapped on each X chromosome (future Xi 129 and future Xa *cast*) by allelic analysis at the XIC (chrX: 103126305-103891745) in D0 cells. The 129 track is in pink (wt) and purple (KO) and the *cast* track in light blue (wt) and dark blue (KO). **(B)** Same analysis as in **(A)**, but zoomed in on the *Xist/Tsix* locus. Gene expression tracks for the 129 and *cast* alleles determined by allelic RNA-seq are shown below. Consistent with previous studies, there is 129-specific (future Xi) enrichment of H3K27me3 across *Xist* [30]. There is no obvious H3K27me3 increase following *Kdm6a* KO. SNPs are shown by gray lines below IGV tracks. The scale of the profiles shown in **(A**-**C)** are indicated in the upper right corners.

## Supplemental tables

1. Summary of clones and primers used
2. DEseq in D0 and D15 *Tsix-*stop cells
3. Germ layer marker gene expression in D0 and D15 *Tsix-*stop cells
4. Genes with increased expression from Xi in differentiated *Kdm6a* KO cells
5. Xi expression of genes in temporally define categories in wt and KO *Tsix*-stop cells
6. RNA-seq allelic mapping metric
7. Allelic H3K27me3 and IgG Cut&Run mapping summary

