## Supplementary figures and images for "KDM6A facilitates Xist upregulation at the onset of X inactivation"

### Supplemental Figures

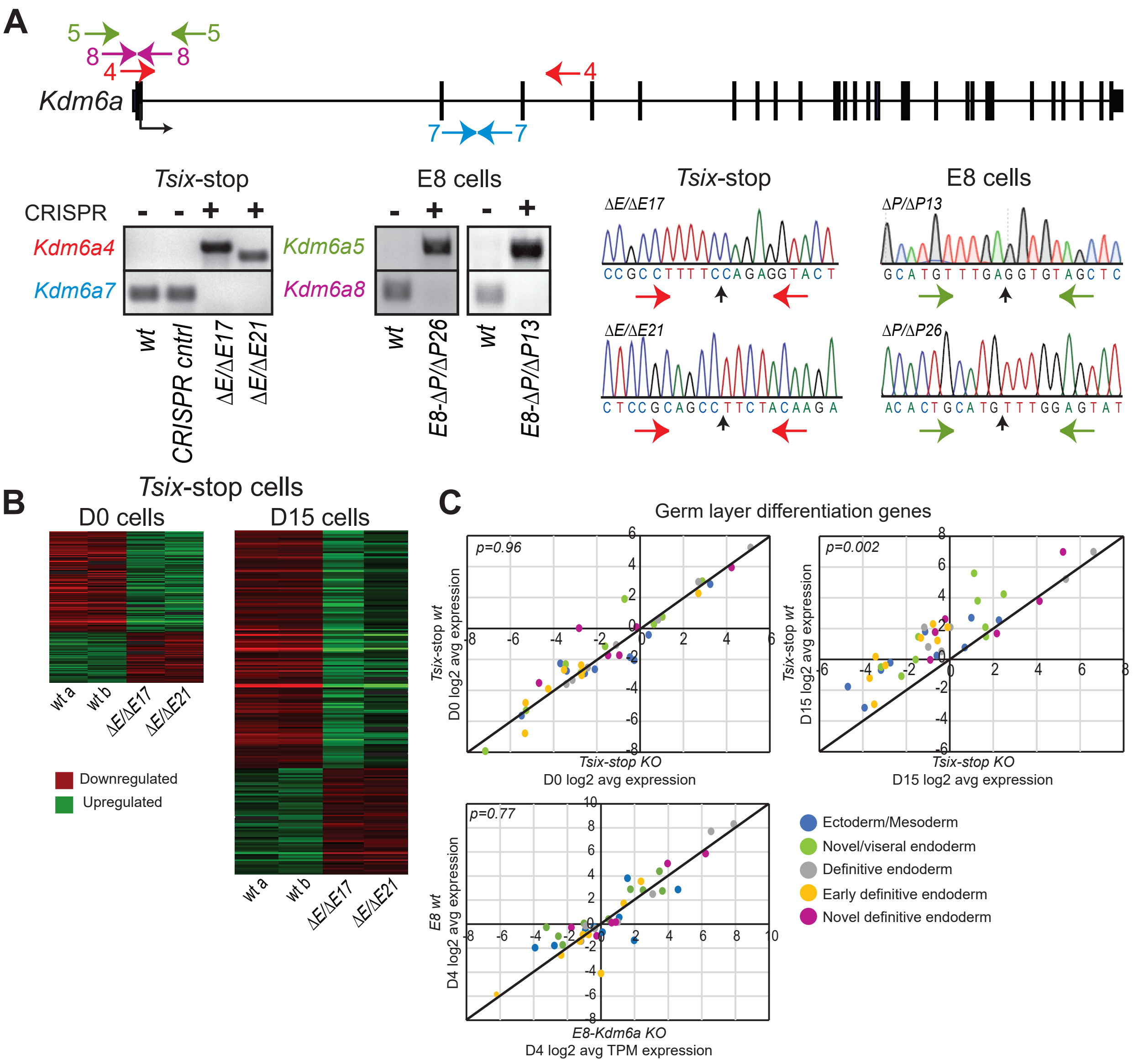

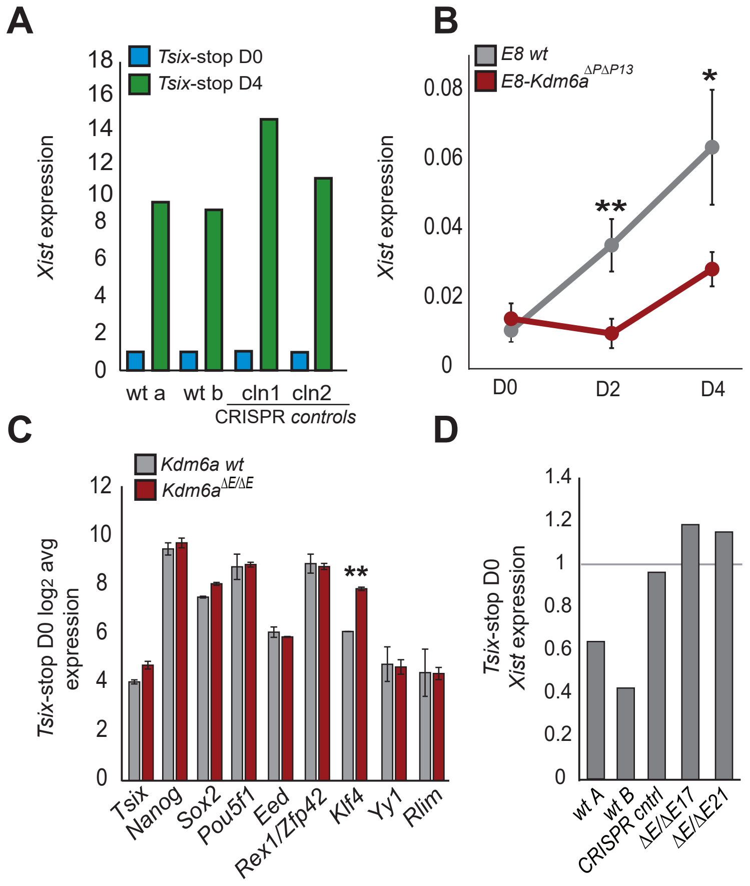

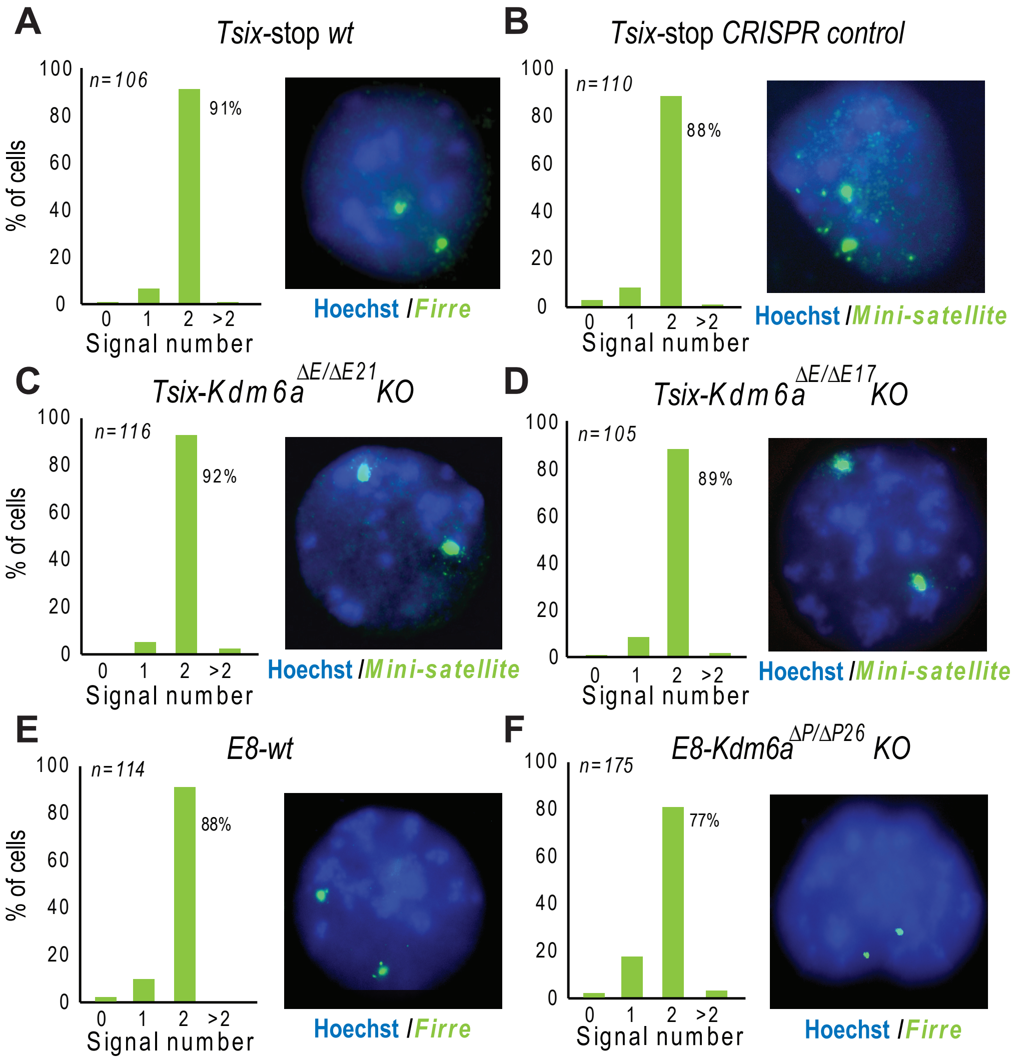

Figure S4

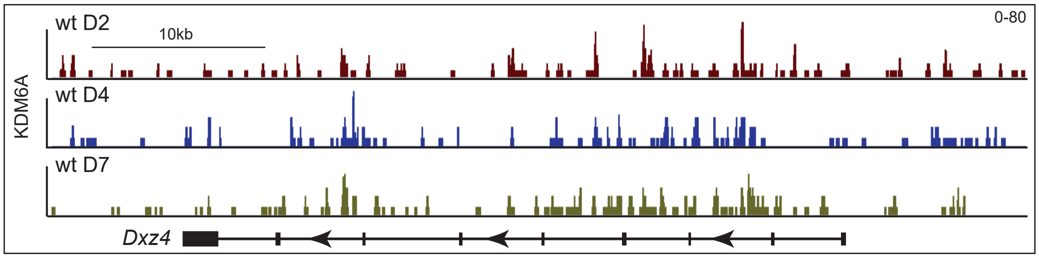

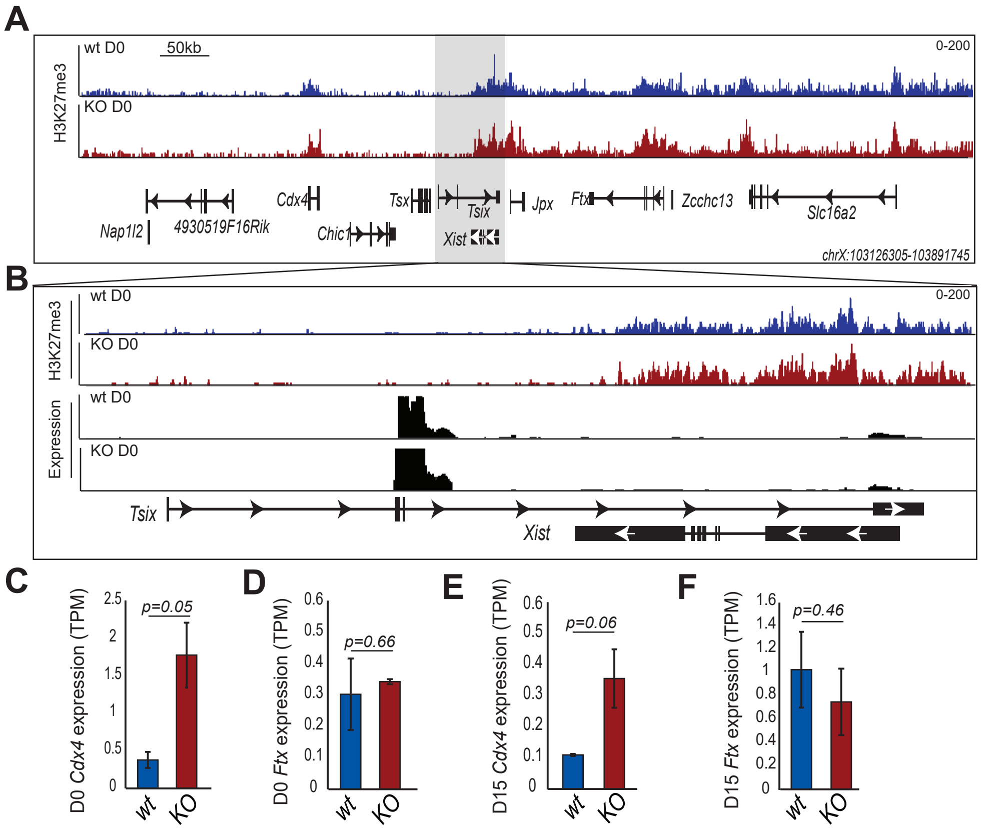

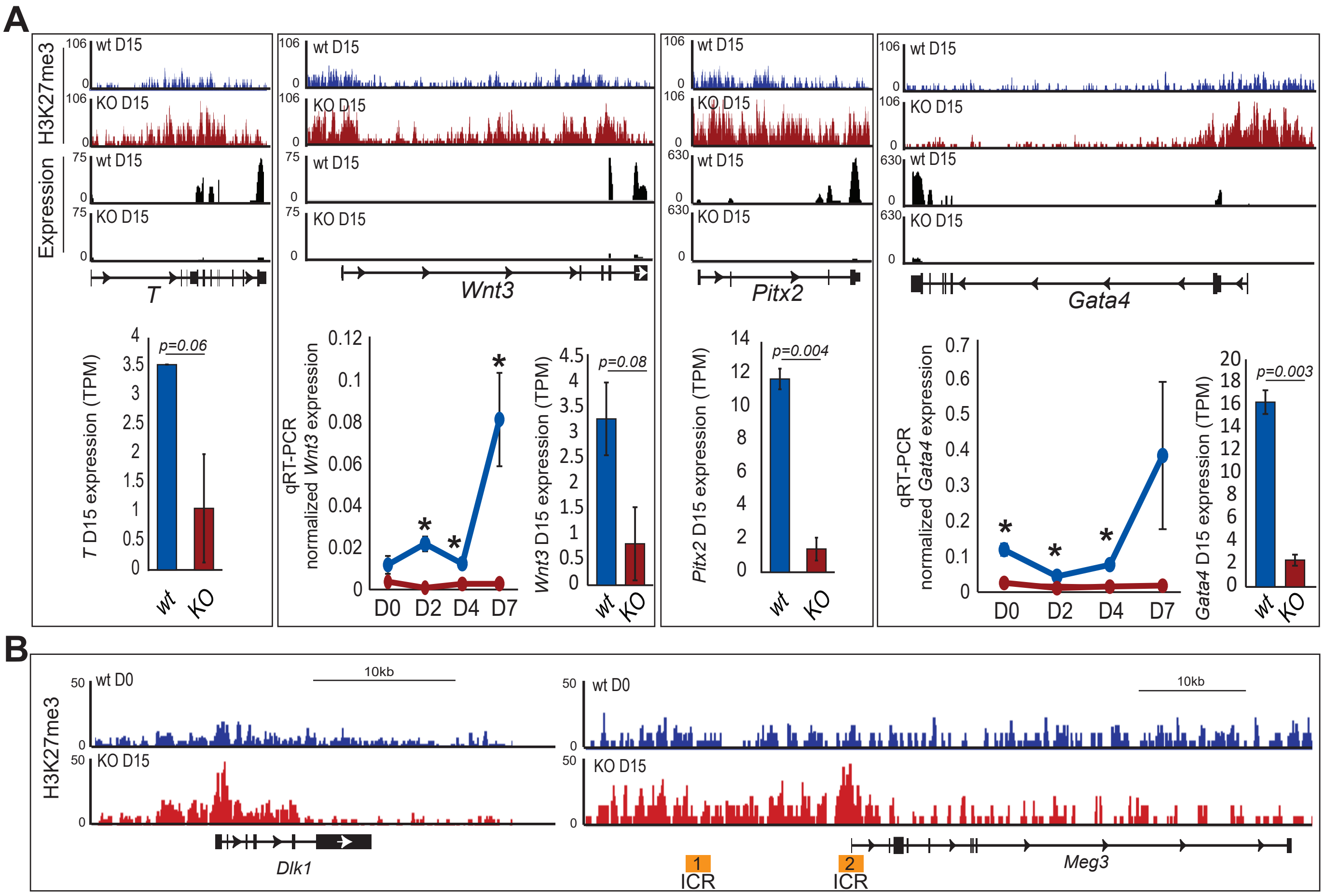

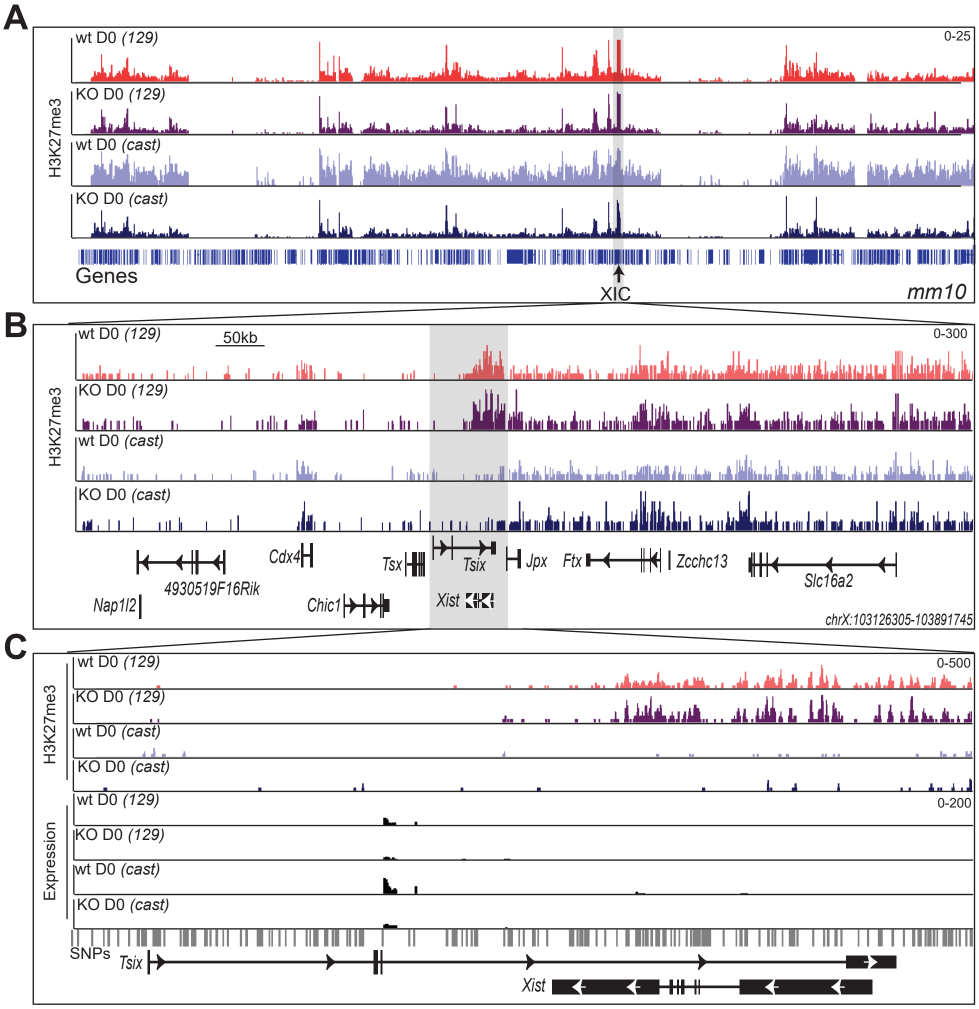
